# TUSCAN: Tumor segmentation and classification analysis in spatial transcriptomics

**DOI:** 10.1101/2024.08.20.608863

**Authors:** Chenxuan Zang, Charles C. Guo, Peng Wei, Ziyi Li

## Abstract

The identification of tumor cells is pivotal to understanding tumor heterogeneity and the tumor microenvironment. Recent advances in spatially resolved transcriptomics (SRT) have revolutionized the way that transcriptomic profiles are characterized and have enabled the simultaneous quantification of transcript locations in intact tissue samples. SRT is a promising alternative method of studying gene expression patterns in spatial domains. Nevertheless, the precise detection of tumor regions within intact tissue remains a great challenge. The common way of identifying tumor cells is via tumor-specific marker gene expression signatures, which is highly dependent on marker accuracy. Another effective approach is through aneuploid copy number events, as most types of cancer exhibit copy number abnormalities. Here, we introduce a novel computational method, called TUSCAN (TUmor Segmentation and Classification ANalysis in spatial transcriptomics), which constructs a spatial copy number variation profile to improve the accuracy of tumor region identification. TUSCAN combines the gene information from SRT data and the hematoxylin-and-eosin-staining image to annotate tumor sections and other benign tissues. We benchmark the performance of TUSCAN and several existing methods through the application to multiple datasets from different SRT platforms. We demonstrate that TUSCAN can effectively delineate tumor regions, with improved accuracy compared to other approaches. Additionally, the output of TUSCAN provides interpretable clonal evolution inferences that may lead to novel insights into disease development and potential druggable targets.

## 1 Introduction

Intratumor heterogeneity, which is characterized by the presence of diverse tumor cell populations, is a driver of disease progression and represents a primary cause of therapeutic resistance. Understanding intratumor heterogeneity and the associated tumor microenvironment is crucial for uncovering the mechanisms behind cancer progression and invasion [1], as well as the modulation of immune cell activities [2]. Such insights not only enhance research into treatment responses [3, 4] but also pave the way for more effective therapeutic interventions. In the past decade, single-cell technology has been successful at providing a comprehensive understanding of cell populations within the tumor microenvironment. Nonetheless, the absence of spatial information limits the ability of single-cell data to discern distinct tumor populations across various tissue locations.

Advances in spatially resolved transcriptomics (SRT) have provided a promising avenue for exploring the spatial distribution of tumor cells [5, 6]. One key advantage of SRT is its ability to preserve cells’ spatial locations during the generation of gene expression profiles. This unique feature enables more profound exploration of gene expression patterns across various tumor regions in a spatial context, thereby allowing for the inference of tumor clonal substructures and tumor evolution analysis. The primary task in these analyses involves accurately pinpointing tumor regions within SRT data.

A conventional way of identifying tumor cells is to use tumor-specific gene markers. For example, previous SRT studies performed initial clustering on SRT spots and defined cluster labels using known cell type markers [7]. A recently developed computational pipeline called TESLA [8] distinguishes tumors from other tissues via a machine learning algorithm and cancer gene markers [9]. However, this approach has several limitations. First, the efficacy of TESLA is contingent upon the selection of specific markers for different cancer types. This reliance introduces significant variability in the results, emphasizing the crucial role of marker accuracy. Second, the identification of consistent and reliable marker genes becomes extremely difficult in highly heterogeneous tumors such as triple-negative breast cancer [10], and melanoma [11], which exhibit significant genomic variability. The tumor cells may even change their phenotype in response to treatment, rendering these markers not universally applicable to all individuals. These circumstances undermine the effectiveness of using typical gene markers for tumor identification.

An alternative approach to identifying tumor regions involves detecting aberrant copy number variations (CNVs), a hallmark that is present in the majority of cancer cells. Although copy number profiles (CNPs) are based on DNA levels [12], several methods, such as inferCNV [13] and CopyKAT [14], have demonstrated the feasibility of inferring CNPs from single-cell RNA sequencing (scRNA-seq) data. These methods leverage the principle that the gene expression levels of adjacent genes along the genome are similar, enabling the estimation of CNPs through the averaged gene expression across genomic loci. Several recent studies have directly applied inferCNV to various spatial transcriptomics cancer datasets [15–17], demonstrating its ability to detect spatial CNV patterns. However, as shown in our experiments, SRT has distinct data characteristics on scRNA-seq data. Directly applying scRNA-seq-based CNV methods filters out important signals in SRT. Additionally, different SRT platforms generate data with distinct features, which should be taken into account in the CNV analysis.

Given these challenges, it is necessary to develop an SRT-specific tool for distinguishing tumor regions from non-tumor tissues while overcoming the limitation of marker selections. Here, we introduce TUSCAN (**TU**mor **S**egmentation and **C**lassification **AN**alysis in spatial transcriptomics), an automated computational pipeline to perform tumor region segmentation and classification analysis by inferring spatial CNPs from gene expression and histology images in spatial transcriptomics data. Inspired by inferCNV [13], our method reconstructs CNPs by comparing the expression levels of genes in each spot to those in diploid spots, using a fixed-size sliding window across the genome to smooth the data. However, unlike inferCNV, our method can automatically select normal reference spots with high confidence using an evaluation score system that combines gene expression and histology images. TUSCAN also tailors the gene filtering, normalization, and normal residual signal neutralization procedures on the basis of the data characteristics from different SRT platforms. We applied TUSCAN to multiple SRT datasets, demonstrating its superior performance on tumor segmentation over existing methods that serve similar purposes. Furthermore, through a case study of human breast cancer SRT data, we illustrated how constructing spatial CNPs can facilitate tumor subclone classification and uncover distinct spatial CNV patterns among tumor subclusters. Moreover, visualization of the CNV signals across spatial locations can aid in exploring cancer progression, understanding tumor lineage relationships, and identifying key mutational events that drive clonal differentiation. All of these efforts offer deeper insight into intra-tumor heterogeneity. TUSCAN is available as user-friendly R software at https://github.com/CZang409/TUSCAN.

## 2 Results

### 2.1 Overview of TUSCAN

TUSCAN is applied to the SRT data generated from sequencing-based platforms such as spatial transcriptomics (ST) [18] and 10x Visium. We assume that both the SRT data (gene expression and location information) and the associated hematoxylin-and-eosin (H&E)-stained images are available for analysis. Figure 1 shows the structured methodology of TUSCAN, which consists of three distinct steps. Step 1: Find a subset of high-quality normal spots. The initial step involves identifying a subset of spots with a high probability of being normal tissue. In order to achieve this goal, we first divide all of the spots into several clusters, followed by the application of a Gaussian mixture model to estimate the gene expression variance within each cluster. Meanwhile, the histology image is transformed into a grayscale plot, from which the gray channel is extracted to compute the mean gray value for each cluster. Step 2: Infer the CNV. The histology image is used to identify normal tissue regions on the basis of two observations: first, the cluster exhibiting the minimum variance in gene expression usually indicates normal tissue, and second, the cluster with the lightest hue in the histology image is more likely to represent normal tissue than those with darker hues. Upon isolating a subset of normal spots, we use the mean gene expression value of these spots as a baseline reference for inferring the CNVs. This genome-wide CNV analysis is performed for all spots. Step 3: Identify the tumor region. The final step of TUSCAN employs consensus clustering to partition the tissue sample into two regions: tumor and normal tissue, predicted on the basis of the derived CNV matrix. A detailed description of these three steps is presented in the Methods section.

**Figure 1:**
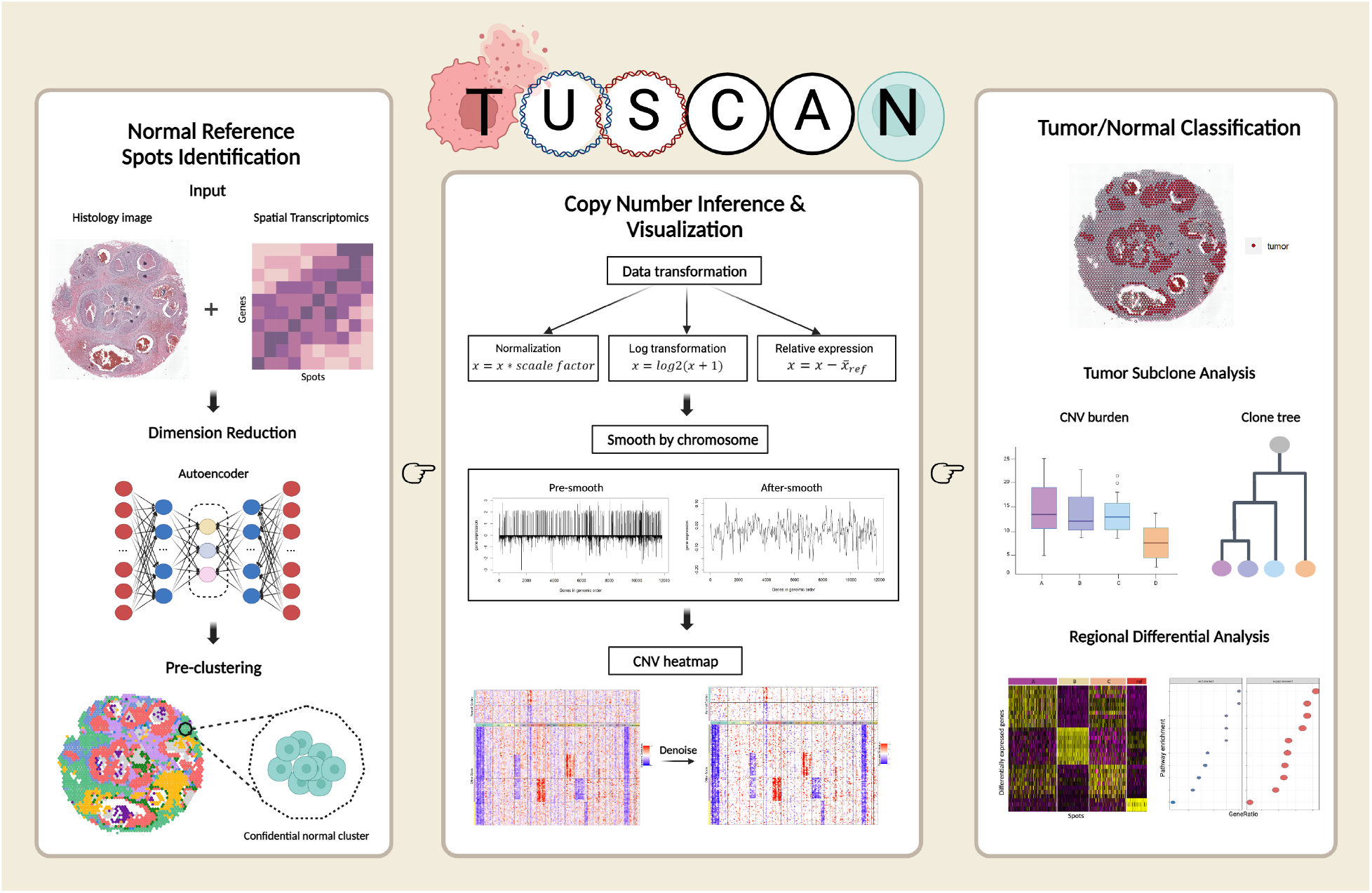
Schematic overview of TUSCAN. TUSCAN is designed to identify tumor regions in spatially resolved transcriptomics data. First, it integrates the histology image and spatial gene expression data as inputs; it then employs an autoencoder for dimensionality reduction. The low-dimensional features serve as the basis for an initial clustering of all spots. The cluster with the highest confidence of representing normal cells is defined (left box). A CNP for all spots is constructed (middle box) with the selected cluster as a normal reference. Finally, TUSCAN segments all spots into tumor regions and normal regions via consensus clustering with the CNP as the input. TUSCAN is also capable of performing a tumor subclone analysis, reconstructing a tumor clonal tree, and performing a regional differential analysis (right box). Created with BioRender.com.

### 2.2 Systematic evaluation of TUSCAN using four SRT datasets

We applied TUSCAN to four publicly available SRT datasets. We studied two human breast cancer datasets from the 10x Visium platform. The first sample was annotated with ductal carcinoma and invasive carcinoma, including 2,518 spots and 17,943 genes (Fig. 2a) [19]. The second sample was annotated with ductal carcinoma in situ, lobular carcinoma in situ and invasive carcinoma, including 3,798 spots and 36,601 genes (Fig. 2b)[20]. To demonstrate the robustness of TUSCAN in different cancer types, we explored another invasive human prostate cancer SRT dataset from the 10x Visium platform, which contains 4,371 spots and 17,943 genes (Fig. 2c) [21]. The last dataset was a HER2-positive tumor sample. The gene expression profile was measured from the ST platform, with 293 spots and 16,148 genes detected (Fig. 2d) [22]. More detailed information about these datasets can be found in Supplementary Table 1. All of the datasets include the pathologist’s annotated H&E-stained images, which are referred to as the ground truth. We compared the tumor domain detection accuracy of TUSCAN with that of three other methods: TESLA [8], CopyKat [14], and BayesSpace [23]. BayesSpace is a widely used Bayesian method of clustering spots in SRT data, offering robust and versatile capabilities for general clustering. We used the Adjusted Rand Index (ARI) [24] to measure the concordance between the identified tumor cluster and the ground truth.

**Figure 2:**
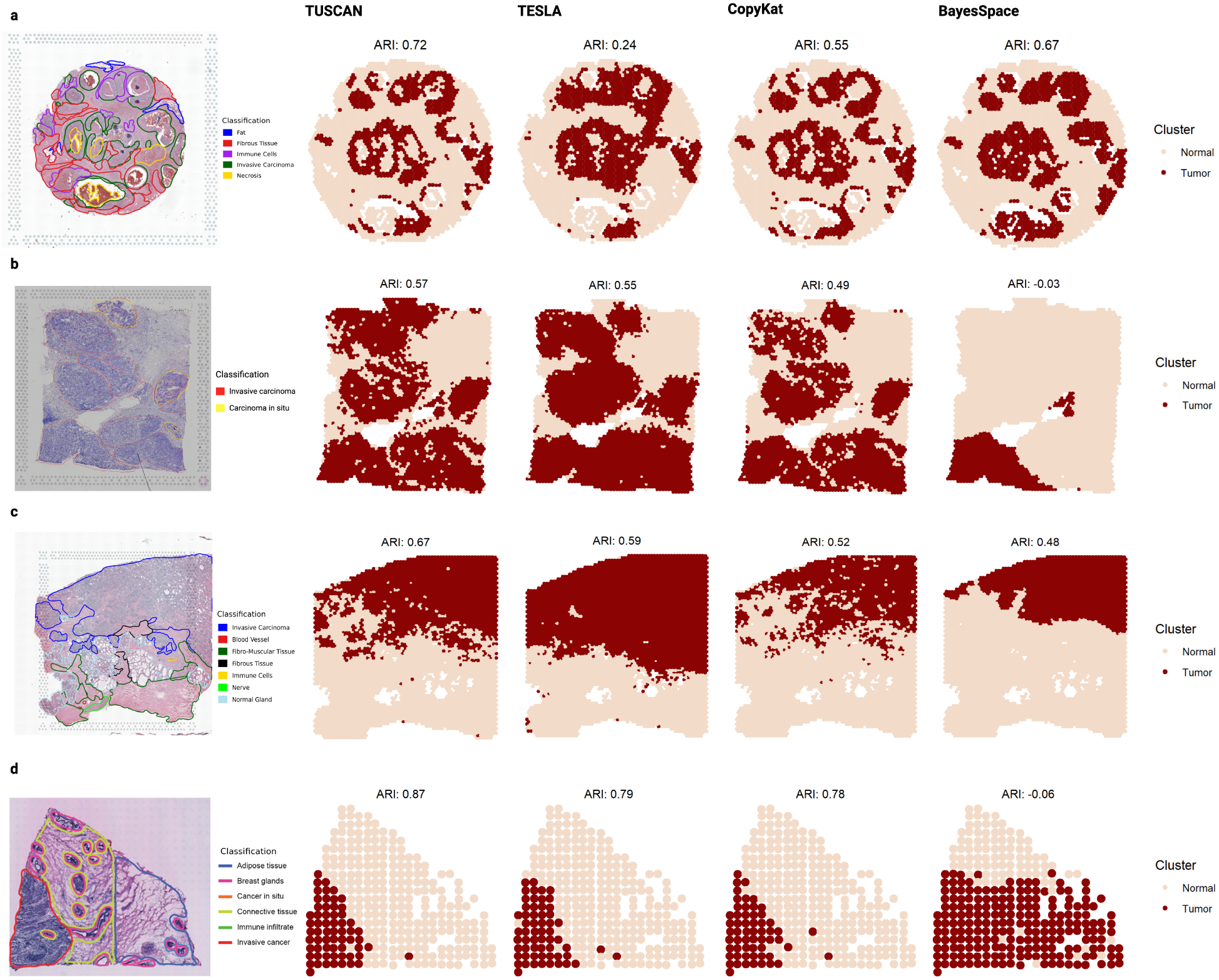
Benchmark evaluation and comparison with other methods. Pathologist-annotated histology images are in the left panel. Tumor sections of four human cancer samples identified by TUSCAN, TESLA, CopyKat, and BayesSpace are marked in red. **a** Human breast cancer (Ductal Carcinoma in Situ). **b** Human breast cancer (Block A, Section 1). **c** Human prostate cancer. **d** Human epidermal growth factor receptor 2+ breast cancer, patient B.

Figure 2 demonstrates that TUSCAN performs best at distinguishing tumor tissues compared with other existing methods, as it achieves the highest ARI values across all datasets. Especially in the prostate cancer dataset, TUSCAN is the only method that results in an ARI value above 0.6. Overall, TESLA is the second-best performing method. However, a significant variation in classification accuracy is evident across three human breast cancer datasets when adopting a uniform set of marker genes to TESLA (Fig. 2a,b,d). Specifically, the analysis of the first dataset is notably inferior to that of the subsequent datasets, indicating an inherent challenge in identifying consistent and reliable markers for cancers that are characterized by high tumor heterogeneity, such as breast cancer, that are universally applicable to different individuals. Furthermore, we noticed that the performance of TESLA is greatly influenced by the choice of its built-in parameter *k*, which denotes the number of genes used to calculate the relative expression and may present an obstacle for end-users. CopyKat also achieves commendable results and demonstrates stable performance across different datasets. It is worth noting that both TUSCAN and CopyKat could distinguish between tumor and necrosis with remarkable precision (Fig. 2a), while BayesSpace indiscriminately categorized necrosis as tumor, and TESLA failed to effectively differentiate between the two morphologies. This demonstrates the advantage of using CNVs in tumor identification. Although CopyKat exhibits a notable true positive rate in tumor segmentation, it has a propensity to miss certain tumor spots. This discrepancy could be attributed to the fact that CopyKat is specifically tailored for high-throughput scRNA-seq data. In contrast, the sequencing depth associated with SRT is generally shallow, further compounding the challenge of accurate tumor spot detection by CopyKat. BayesSpace broadly has the worse performance here, demonstrating that general-purposed clustering methods may lead to biased tumor identification results.

A noteworthy strength of TUSCAN is its ability to precisely delineate tumor boundaries. For example, in the human prostate cancer tissue, the tumor region is clearly composed of a predominant tumor mass accompanied by several smaller adjacent tumors. TUSCAN successfully outlines these tumors, whereas TESLA mixes them together (Figure 2c).

### 2.3 Case study: Human Breast Cancer (Block A, Section 1)

We conducted a comprehensive case study of the human breast cancer (Block A, Section 1) dataset [20]. This sampled tissue was classified as AJCC/UICC Stage Group IIA, with positive estrogen receptor status, negative progesterone receptor status, and positive human epidermal growth factor receptor 2 status. Ductal carcinoma in situ, lobular carcinoma in situ, and invasive carcinoma were present within the sections examined. We first undertook a subclone analysis of the tumor region based on the CNPs computed by TUSCAN. We also constructed a tumor clonal tree to study the tumor evolution process and tumor heterogeneity. Furthermore, we performed a differentially expressed gene (DEG) analysis and gene set enrichment analysis (GSEA) to explore the distinct patterns among tumor subclones.

#### 2.3.1 Clone substructure heterogeneity analysis

The copy number matrix inferred from TUSCAN was extracted and clustered to identify potential tumor subpopulations. 2265 tumor spots discerned by TUSCAN were divided into three main subclones (clones 1, 2, and 3). In Figure 3a, we illustrate the spatial distribution of these distinct tumor clones, mapped onto a histology image of the intact tissue. Each clone is confined to a well-defined region, indicating non-random spatial patterns with minimal intermingling between the subpopulations. The heatmap (Fig. 3b) demonstrates that the three clones share certain similar CNV patterns, including copy number gains (1q, 8q, 17q) and copy number losses (7q, 11q). These genomic regions included many known breast cancer genes, for example, *MDM4, MYC, CCND1, ERBB2*, and *BRCA1* [25]. In addition to the common CNVs, each clone exhibits its distinct pattern. For example, clone 1 is characterized by a larger number of amplifications (3p, 5q, 12q, 16p, 20q) than clones 2 and 3 (Fig. 3b,c). Clone 2 reveals increased amplifications on 1q and additional deletions on 17p and 22q. Clone 3 exhibits the most complex CNV patterns, which are discussed in more details in the next section.

**Figure 3:**
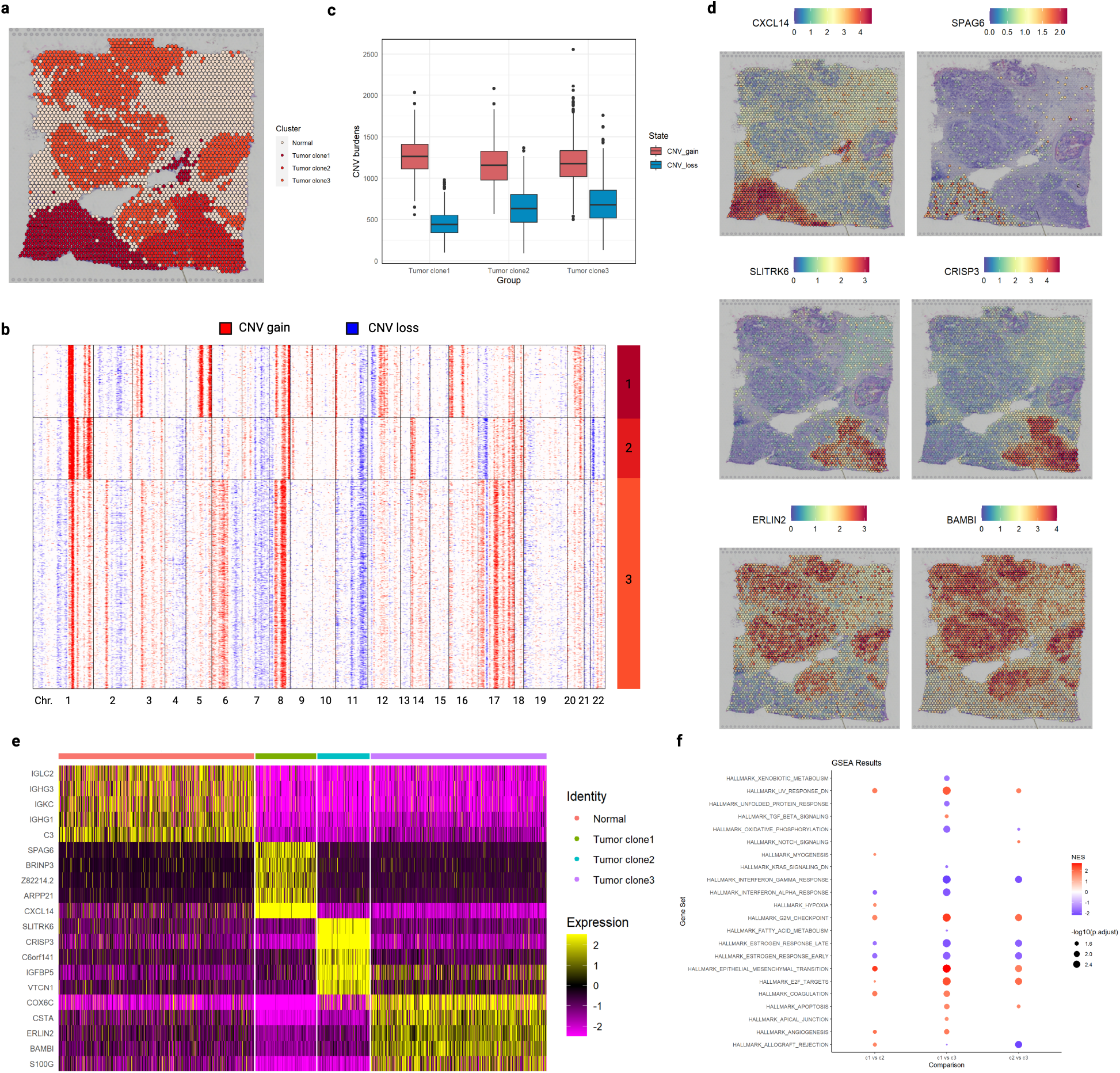
Tumor clone analysis of human breast cancer. **a** Spatial visualization of three tumor clones and normal tissue sections. **b** Heatmap of the estimated CNP of tumor spots. The clonal classifications of spots were determined by consensus clustering. **c** CNV burden is calculated by the number of genes shown at extreme expression levels compared to the normal group. CNV_gain: *e*_*i*_ *> mean*(*e*_*i normal*_)+2*\*sd*(*e*_*i normal*_). CNV loss: *e*_*i*_ *< mean*(*e*_*i normal*_) *−* 2 *\* sd*(*e*_*i normal*_). **d** Spatially variable features across three tumor clones. Top to bottom: clone 1 vs. clones 2 and 3; clone 2 vs. clones 1 and 3; clone 3 vs. clones 1 and 2. **e** Differential expression analysis of three tumor clones and normal cells. **f** GSEA of three tumor clones. Human hallmark gene sets from Molecular Signature Database were used as the reference source.

To further study the tumor heterogeneity, we conducted a DEG analysis and GSEA across these three tumor clones (Fig. 3d-f). Clone 1 had a significant increase in the expression levels of several genes that are involved in inflammatory responses and the regulation of the cell cycle, such as *CXCL14* and *CCND1* [26, 27]. For clone 2, there was a notable elevation in *CRISP3*, a gene that is closely linked to the immune response, indicating that it has a significant role in this clone’s distinct genetic profile [28]. Our analysis of clone 3 highlighted an enhanced expression of key genes such as *CSTA, ERLIN2*, and *S100G*, which are crucial for preserving both the structural and functional integrity of cellular components [29–31].

On the basis of the GSEA results (Fig. 3f), clone 1 demonstrates significant enrichment of the genes that define epithelial-mesenchymal transition, genes that are down-regulated in response to ultraviolet radiation, and genes that are associated with blood coagulation compared to clones 2 and 3. Epithelial-mesenchymal transition is closely related to cellular motility. Cells that are enriched in this pathway may have enhanced migratory and invasive capabilities, suggesting that clone 1 possesses higher aggressiveness [32, 33]. It has a reduced ability to respond to ultraviolet damage, indicating abnormalities in the DNA damage repair pathways [34–36] and potentially leading to accumulated mutation events and tumor progression. Moreover, coagulation is positively correlated with angiogenesis and might play an important role in the tumor microenvironment [37].

On the other hand, tumor clone 1 shows suppression of interferon alpha and gamma responses, suggesting potential downregulation of immune response signaling compared to clone 3 [38–40]. In addition, genes that are involved in both the late and early estrogen responses are down-regulated in clones 1 and 2 compared to in clone 3, with clone 1 showing more significant changes. The GSEA results suggest that tumor clone 1 exhibits a higher degree of malignancy and enhanced invasive capabilities compared to clones 2 and 3.

#### 2.3.2 Tumor clonal tree reconstruction

On the basis of the observations shown in Figures 3a and 3b, tumor clone 3 occupies the majority of the tissue sample and exhibits a complex CNV pattern. Consequently, we further divided clone 3 into four subgroups according to its CNV profile, along with clones 1 and 2, and designated these as clones A through F (Fig. 4a). Figure 4b presents the heatmap of CNPs for these six tumor clones. It is clearly shown that although clones C to F (clone 3) share similar CNV patterns, distinct differences are also present. We then constructed a phylogenetic tree, presented in Figure 4c to illustrate the evolutionary trajectories of the six tumor clones on the basis of their CNV profiles, with gray nodes and one pink node representing the inferred unknown ancestral clones and original normal cells, respectively. Clones A and B exhibit distinct evolutionary paths as shown by the divergent branches originating from the normal cells, which may indicate that clones A and B may not originate from the same common clone as clones C to F. Clone D is the root within clone 3, demonstrating its status as an earlier-formed population within this cluster. Upon referring to the pathologist’s annotations, it becomes evident that clone D essentially covers the carcinoma in situ region. The high consistency between clone D and the pathologist-delineated carcinoma in situ area highlights the advantages of employing clonal for mapping tumor progression and identifying unique tumor subregions. We performed trajectory analysis on tumor regions on the basis of CNPs generated from TUSCAN. The inferred pseudo-times indicates that clone 3 was formed earliest, while clones 2 and 3 were formed at a later stage (Fig. 4d). These findings facilitate the further detailed examination of CNV loci within each subclone, enabling the discovery of distinct CNV patterns and the identification of specific gene markers across different tumor subclones.

**Figure 4:**
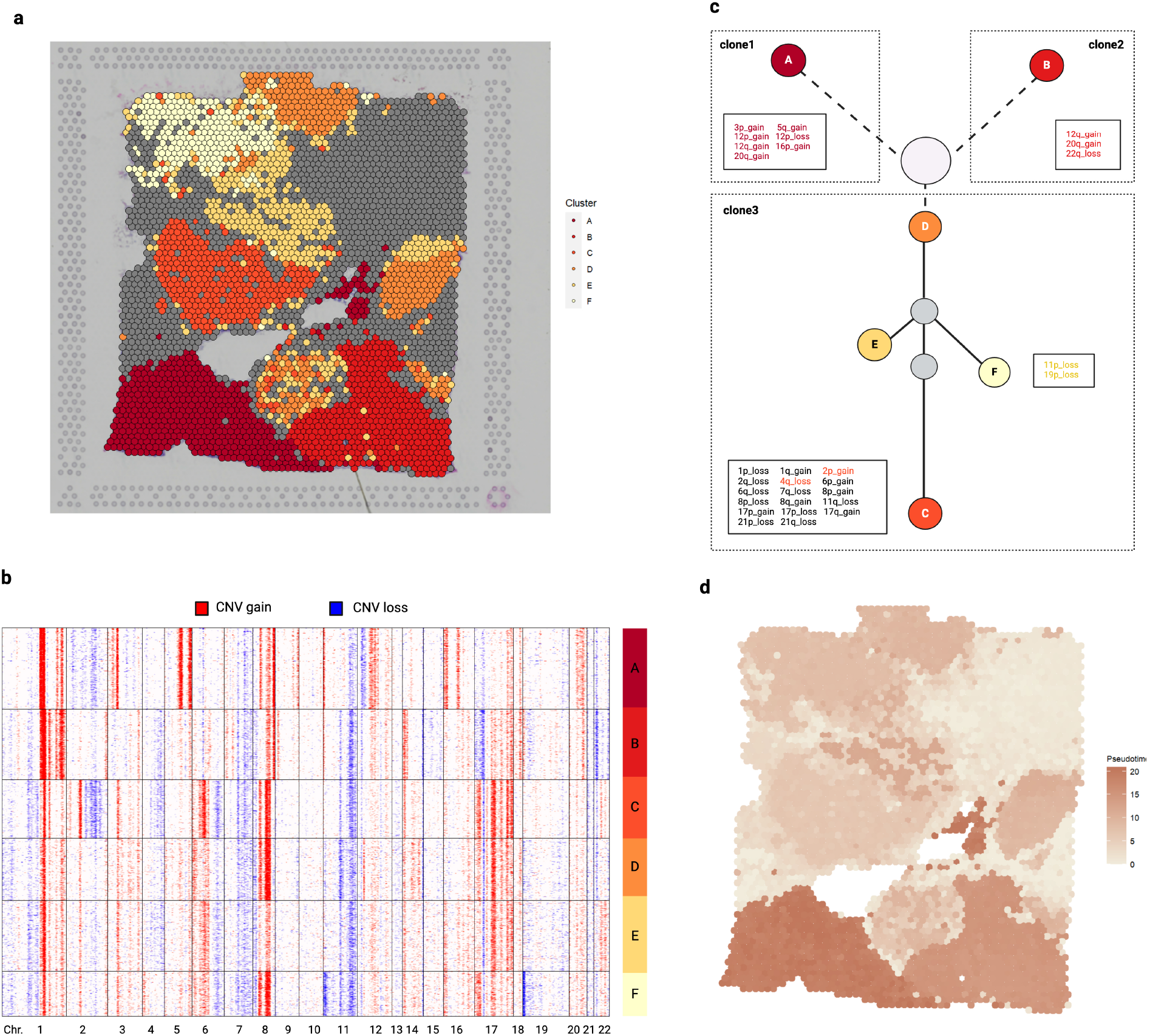
Further sub-clone tree reconstruction. **a** Spatial visualization of six tumor clones. **b** Heatmap of estimated CNPs for tumor spots. Clone 3 was further separated into four subclones. **c** Phylogenetic clonal tree of the tumor clones from inferred CNPs. Gray circles are unknown ancestors. The pink circle represents the original normal cells. All CNV locations of clone C are marked on the left side of the node, where the maximum CNV locations are observed. CNV locations that are unique to other clones in comparison to clone C are indicated with distinct colors. The complete CNV profile can be found in the Supplementary Materials Table S3. **d** Visualization of the inferred trajectory. Higher pseudo-times are marked as darker colors.

## 3 Discussion

In this paper, we presented TUSCAN, a statistical method that integrates gene expression data and histology images to infer spatial CNVs for tumor region segmentation. Using four different real cancer dataset analyses, we demonstrated that TUSCAN had higher accuracy at identifying tumor domains than existing methods. A notable strength of our approach lies in its capability to autonomously delineate tumor regions, making it broadly applicable across a diverse array of cancer types. This characteristic obviates the necessity for cancer-specific marker genes, thereby rendering our method particularly advantageous for those cancers that do not have established gene markers. The universality of TUSCAN not only facilitates its application but also significantly improves user accessibility and implementation across different cancer research contexts, aligning with the interdisciplinary nature of modern oncological studies.

By constructing spatial CNV profiles, we have been able to distinctly delineate copy number amplifications and deletions within specific chromosomal bands in cancer samples. This allows for a deeper understanding of intratumoral heterogeneity and aids in the delineation of tumor subclones. In our breast cancer case study, we discerned three major subclusters (clone 1 to 3), which were further subdivided into six distinct subclones (clones A to F). Through the construction of a clonal tree complemented by a trajectory analysis, we were able to delineate the evolutionary trajectory of the tumor. Our findings suggest that subclones C to F, which constitute clone 3, likely emerged at an early stage in the tumor’s development. In contrast, subclones A and B (or clones 1 and 2) appeared at a later phase. Moreover, our comprehensive DEG analysis and GSEA results elucidated significant variations in gene signatures and pathways across the different tumor substructures. Notably, these analyses highlighted disparities in critical biological processes and pathways such as epithelial-mesenchymal transition, response to ultraviolet damage, coagulation, cellular responses to alpha and gamma stimuli, and apoptosis.

We acknowledge several limitations of TUSCAN. First, owing to the inherent spot-based design of the technology, wherein each spot includes multiple cells (e.g., ST and 10x Visium), TUSCAN cannot achieve single-cell resolution. Consequently, the tumor spots identified through our method may not possess absolute purity. Second, TUSCAN might not be a suitable choice for analyzing cancer types where CNV events are rare or absent, such as chronic myeloid leukemia and acute myeloid leukemia [41, 42]. Third, given the necessity of inferring the CNPs from the entire transcriptome, TUSCAN is not applicable to imaging-based SRT technologies, for example, MERFISH and seqFISH, which are capable of simultaneously measuring only hundreds of genes. Our future work will be focused on integrating multi-omics data to refine resolution and augment the precise tumor identification. Moreover, we plan to expand our efforts towards the annotation of a broader spectrum of cell types, enriching the depth and breadth of our analyses.

## 4 Methods

### 4.1 Tumor segmentation

#### Step 1: Find a subset of high-quality normal cells

The overall workflow of TUSCAN is shown in Figure 1. To accurately infer CNV through SRT data, we first need to recognize a subset of normal spots with high confidence as the reference baseline values. We conduct a preliminary clustering on the basis of the spatial gene expression data and the H&E-stained image. The gene expression data is denoted by an *N × D* matrix of unique molecular identifiers where *N* represents genes and *D* represents spots. The raw gene expression matrix is normalized and log-transformed through the standard approach in the single-cell RNA data processing procedure. We then employ an autoencoder on the top 2000 highly variable genes to perform dimension reduction. By default, we obtain 20 dimensions, but users can also customize the number of dimensions after reduction. In addition to gene expression, we believe that the H&E-stained histology image provides invaluable insights for spatial clustering. Thus, for each spot *i*, we extract the three-channel RGB values by mapping each spot to the original histology image according to its 2D pixel coordinates (*x*_*i*_, *y*_*i*_). However, since the number of pixels is much larger than that of spots, using the RGB values of a single pixel to represent each spot may cause a significant bias. Therefore, we draw a 99×99 square centered on (*x*_*i*_, *y*_*i*_) and apply the average of the color within the square as the RGB (*r*_*i*_, *g*_*i*_, *b*_*i*_) values for spot *i*. Next, three-color values are scaled to the same range as the 20 components, and these 23 features are used for preliminary clustering (Louvain or K-means). We recommend increasing the number of clusters while ensuring that the reference cluster contains at least 10 spots to reduce bias.

To determine the cluster with the highest potential purity of normal tissues, we create an evaluation score *s*_*c*_ that combines the gene expression and histology image information as:

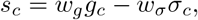

where *g*_*c*_ is the median gray value of cluster *c* (*g*_*i*_ = 0.299 *× r*_*i*_ + 0.587 *× g*_*i*_ + 0.114 *× b*_*i*_, for each spot *i*) and *σ*_*c*_ is the standard deviation of gene expression within cluster *c*, estimated by the gaussian mixture model.

To avoid one feature dominating the other, both *g*_*c*_ and *σ*_*c*_ are standardized. *w*_*g*_ and *w*_*σ*_ are the weights of *g*_*c*_ and *σ*_*c*_, respectively. By default, the weights of gene expression and histology image are set to be equal, which means *w*_*g*_ = *w*_*σ*_ = 0.5. We select the cluster with the maximum *s*_*c*_ as the reference group on the basis of two observations. First, the cluster with the lightest color on the histology image is most likely to represent normal tissue. The rationale is that tumor cells may exhibit larger and more intensely stained nuclei due to an increased nuclear-cytoplasmic ratio, thus appearing darker than normal cells in the H&E-stained image. Second, the cluster with the lowest variance in gene expression is likely to represent normal tissue.

#### Step 2: Infer the CNP

Intuitively, this step calculates a corrected moving average of gene expression data to determine CNV profiles. First, genes are sorted by absolute genomic position. Specifically, they are first ordered by chromosome and then by genomic start position within the chromosome. The underlying reason for this algorithm is that averaging the expression of genomically adjacent genes removes gene-specific expression variability and yields profiles that reflect chromosomal CNVs. To further refine the CNV profile of tumor cells, we construct the CNV profile of a known normal sample, and then for each gene and cell, the normal sample is subtracted from the tumor sample to determine the final tumor CNV profile.

#### Data preprocessing and transformation

We first filter out those genes that have an average expression level of less than 0.01 in each spot and those that are detected in fewer than three spots. In the next step, we normalize the raw gene count matrix for each spot by the sum of unique molecular identifiers counts and multiply it by a scale factor (median unique molecular identifiers counts per spot). If the data are generated from the ST platform, we skip this step, as ST has a relatively low sequencing depth, and normalization would attenuate the CNV signal. We then perform a standard log-transformation on the data: *x* = *log*_2_(*x* + 1).

#### Obtain the relative gene expression values

The gene expression values of all spots are subtracted by the average value of the normal reference. Since the data have been log-transformed, the resulting values represent the log-fold change differences relative to the control group. The outliers are handled as follows:

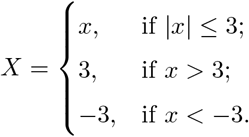

#### Smooth along chromosome

For a given gene *j*, the smoothed expression value *x*_*j*_ is calculated as the weighted average of the genes within a window size *d*:

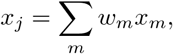

where 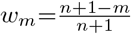 is the weight for the *m*^*th*^ gene near both sides of gene *j, n* = (*d −* 1)*/*2. The gene closer to the center gene *j* is assigned a larger weight. By default, we choose the window size *d* = 101.

#### Further refine the CNV profile

In the next step, the gene expression of each spot is centered by subtracting its median expression intensity. The average expression of the normal reference is subtracted again to further compensate for differences accrued after the smoothing process. The log transformation is reversed to symmetrically represent CNV gain or loss compared to neutral status.

#### Reduce noise

To highlight the CNV signals of tumor, we replace all gene values within 1.5 standard deviations of the average level in normal tissues with the mean expression values of normal tissues.

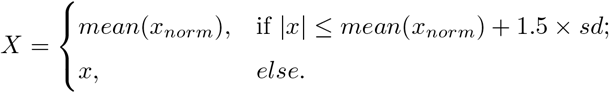

#### Step 3: identify the tumor region

We use a consensus clustering algorithm implemented in the R package `**ConsensusClusterPlus**`[43], to classify all spots into two clusters, tumor and normal, using the copy number matrix obtained from step 2 as input. The process is initiated by selecting a subset of both spots and genes from the copy number matrix. Each subset undergoes clustering *k* groups (*k* = 2 in our case), using a clustering algorithm such as hierarchical clustering, K-means, or a customized method. This clustering step is repeated multiple times.

Then pairwise consensus values, which is the frequency of two items being clustered together across various runs, will be computed to construct a consensus matrix. The algorithm then performs a final round of agglomerative hierarchical consensus clustering using a distance measure based on 1 minus the consensus values, referred to as the consensus clusters. Finally, the distance is calculated between the centroids of two groups and that of the reference group. The group with the larger distance is marked as the tumor.

We evaluate the performance of tumor segmentation using the adjusted rand index (ARI) [24]. The ARI formula is defined as:

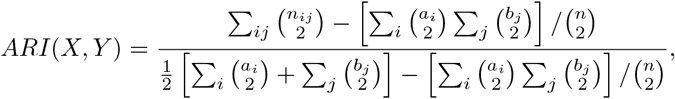

where *X* = *{x*_1_, *x*_2_, …, *x*_*i*_*}* is our spot label, *Y* = *{y*_1_, *y*_2_, …, *y*_*j*_*}* is the true spot label, *n* is the total number of spots, *n*_*ij*_ is the number of spots in both cluster *X* and cluster *Y*, *a*_*i*_ is the total number of spots assigned in *x*_*i*_, and *b*_*j*_ is the total number of spots assigned in *y*_*j*_. The range of ARI is from -1 to 1, with a higher ARI value indicating a better clustering result.

### 4.2 Implementation details of existing methods

In the application of TESLA to the analysis of three human breast cancer datasets, we used 12 marker genes as the meta gene for breast cancer provided in the original TESLA publication: *ERBB2, CNN1, CDH1, KRT5, KRT7, KRT14, KRT18, CDNND1, GATA3, FOXA1, PIP*, and *SCGB2A2* [8]. For the human prostate cancer dataset, nine marker genes were selected from the existing literature to serve as the meta gene: *ADGRF1, CRISP3, TMEFF2, KCNC2, EGR, AMACR, OR51C1P, OR51E2*, and *MYO6* [15,44]. TESLA also requires an input parameter ‘*k*’ to calculate the meta gene’s relative expression at each superpixel, which is defined as the minimum value of the top *k* expressed genes from the meta gene list. Given that TESLA does not suggest how to determine the optimal value of *k*, we applied it with a range of *k* values to ascertain the most effective parameter. CopyKat was executed using the default settings. The classification results were obtained from the prediction result file, where ‘aneuploid’ was marked as tumor. In the case of BayesSpace, due to its inability to directly label the tumor and other tissues, we first employed it to classify all spots into two groups, which were subsequently labeled on the basis of pathologist’s annotations. All other parameters were set to their default values.

### 4.3 Tumor clone tree construction

Our tumor clone tree is constructed on the basis of the maximum-parsimony clone tree computed by the R package `**phangorn**`[45]. The ‘user-defined input’ matrix to the R package phangorn is given in Supplementary Materials Table S2. A detailed tutorial can be found in their vignettes titled ‘Ancestral Sequence Reconstruction’. For the clone tree, the diameter of each node is proportional to the proportion of spots for this clone in the entire sample. The branch lengths are extracted from the tree object created by `**phangorn**`.

### 4.4 Trajectory inference

As illustrated in Figure 4A, the tumor section is partitioned into six sub-clones, labeled A through F. We aim to delve deeper into the developmental trajectory of these tumor clones. For this purpose, we apply the R package `**Slingshot**`[46] to estimate the pseudotime associated with each individual spot. The CNV matrix serves as the initial input for this analysis, upon which UMAP dimensionality reduction is subsequently applied. Before the trajectory analysis, the normal cluster is chosen as the start cluster. The Pseudotime for each spot is extracted using the `**slingPseudotime()**`function and then projected onto the histology image, facilitating a comprehensive visualization of the tumor’s evolutionary dynamics.

### 4.5 DEG analysis and GSEA

A DEG analysis of the human breast cancer data is performed using `**Seurat v4**`. The significantly up regulated genes in each tumor clone compared to other clones and normal tissues are identified by the `**FindMarkers()**`function, with an adjusted p-value of 0.05. We further conduct a pairwise DEG analysis across all clones and perform GSEA on the detected DEGs using the `**GSEA()**`function in `**clusterProfiler**`[47] package employing default settings and an adjusted Benjamini-Hochberg p-value of 0.05. Human hallmark gene sets downloaded from the Molecular Signature Database are used as the reference map.

## Acknowledgments

This work was supported by the Cancer Prevention and Research Institute of Texas (CPRIT) Grant RP230166. We acknowledge the editing services provided by the Research Medical Library at The University of Texas MD Anderson Cancer Center.

## Author Contributions

Z.L. and P.W. conceived the idea and designed the methods. C.Z. conducted the experiments with inputs from Z.L. and P.W. C.G. assisted with obtaining pathologist-annotated cell-type labels and interpreting results. All authors contributed to the writing of the manuscript.

## Data Availability

We analyzed four publicly available SRT datasets: (1) Human Breast Cancer (Ductal Carcinoma In Situ) 10x Visium data; (2) Human Breast Cancer (Block A, Section 1) 10x Visium data; (3) Human Prostate Cancer 10x Visium data; and (4) Human HER-positive breast tumors ST data. The 10x Visium data can be accessed from the 10x Genomics website: https://support.10xgenomics.com/spatial-gene-expression/datasets. Human HER-positive breast tumors ST data can be accessible from their github: https://github.com/almaan/her2st. The details of these datasets are described in Supplementary Table 1.

## Code availability

An open-source implementation of the TUSCAN algorithm in R is available at https://github.com/CZang409/TUSCAN

## References

[1] Q. Wang et al., “Role of tumor microenvironment in cancer progression and therapeutic strategy,” Cancer Medicine, vol. 12, no. 10, pp. 11 149–11 165, 2023.

[2] T. L. Whiteside, “The role of immune cells in the tumor microenvironment,” The Link Between Inflammation and Cancer: Wounds That Do Not Heal, pp. 103–124, 2006.

[3] C. M. Neophytou, M. Panagi, T. Stylianopoulos, and P. Papageorgis, “The role of tumor microenvironment in cancer metastasis: Molecular mechanisms and therapeutic opportunities,” Cancers, vol. 13, no. 9, p. 2053, 2021.

[4] Y. Xiao and D. Yu, “Tumor microenvironment as a therapeutic target in cancer,” Pharmacology & therapeutics, vol. 221, p. 107 753, 2021.

[5] L. Moses and L. Pachter, “Museum of spatial transcriptomics,” Nature methods, vol. 19, no. 5, pp. 534– 546, 2022.

[6] C. G. Williams, H. J. Lee, T. Asatsuma, R. Vento-Tormo, and A. Haque, “An introduction to spatial transcriptomics for biomedical research,” Genome Medicine, vol. 14, no. 1, p. 68, 2022.

[7] S. Z. Wu et al., “A single-cell and spatially resolved atlas of human breast cancers,” Nature genetics, vol. 53, no. 9, pp. 1334–1347, 2021.

[8] J. Hu et al., “Deciphering tumor ecosystems at super resolution from spatial transcriptomics with tesla,” Cell systems, vol. 14, no. 5, pp. 404–417, 2023.

[9] A. J. Bosma et al., “Detection of circulating breast tumor cells by differential expression of marker genes,” Clinical cancer research, vol. 8, no. 6, pp. 1871–1877, 2002.

[10] K. Asleh, N. Riaz, and T. O. Nielsen, “Heterogeneity of triple negative breast cancer: Current advances in subtyping and treatment implications,” Journal of Experimental & Clinical Cancer Research, vol. 41, no. 1, p. 265, 2022.

[11] S. M. Hossain and M. R. Eccles, “Phenotype switching and the melanoma microenvironment; impact on immunotherapy and drug resistance,” International Journal of Molecular Sciences, vol. 24, no. 2, p. 1601, 2023.

[12] A. B. Olshen, E. S. Venkatraman, R. Lucito, and M. Wigler, “Circular binary segmentation for the analysis of array-based dna copy number data,” Biostatistics, vol. 5, no. 4, pp. 557–572, 2004.

[13] A. P. Patel et al., “Single-cell rna-seq highlights intratumoral heterogeneity in primary glioblastoma,” Science, vol. 344, no. 6190, pp. 1396–1401, 2014.

[14] R. Gao et al., “Delineating copy number and clonal substructure in human tumors from single-cell transcriptomes,” Nature biotechnology, vol. 39, no. 5, pp. 599–608, 2021.

[15] A. Erickson et al., “Spatially resolved clonal copy number alterations in benign and malignant tissue,” Nature, vol. 608, no. 7922, pp. 360–367, 2022.

[16] E. Denisenko et al., “Spatial transcriptomics reveals ovarian cancer subclones with distinct tumour microenvironments,” bioRxiv, pp. 2022–08, 2022.

[17] L. Chen et al., “Visualizing somatic alterations in spatial transcriptomics data of skin cancer,” bioRxiv, pp. 2022–12, 2022.

[18] P. L. Ståhl et al., “Visualization and analysis of gene expression in tissue sections by spatial transcriptomics,” Science, vol. 353, no. 6294, pp. 78–82, 2016.

[19] “Human breast cancer (ductal carcinoma in situ),” 2021. [Online]. Available: https://www.10xgenomics.com/datasets/human-breast-cancer-ductal-carcinoma-in-situ-invasive-carcinoma-ffpe-1-standard-1-3-0.

[20] “Human breast cancer (block a section 1),” 2020. [Online]. Available: https://support.10xgenomics.com/spatial-gene-expression/datasets/1.1.0/V1_Breast_Cancer_Block_A_Section_1.

[21] “Human prostate cancer,”2021. [Online]. Available: https://www.10xgenomics.com/datasets/human-prostate-cancer-adenocarcinoma-with-invasive-carcinoma-ffpe-1-standard-1-3-0.

[22] A. Andersson et al., “Spatial deconvolution of her2-positive breast cancer delineates tumor-associated cell type interactions,” Nature communications, vol. 12, no. 1, p. 6012, 2021.

[23] E. Zhao et al., “Spatial transcriptomics at subspot resolution with bayesspace,” Nature biotechnology, vol. 39, no. 11, pp. 1375–1384, 2021.

[24] W. M. Rand, “Objective criteria for the evaluation of clustering methods,” Journal of the American Statistical association, vol. 66, no. 336, pp. 846–850, 1971.

[25] C. G. A. Network, “Comprehensive molecular portraits of human breast tumours,” Nature, vol. 490, no. 7418, pp. 61–70, 2012.

[26] S. Elsheikh et al., “Ccnd1 amplification and cyclin d1 expression in breast cancer and their relation with proteomic subgroups and patient outcome,” Breast cancer research and treatment, vol. 109, pp. 325– 335, 2008.

[27] J. Lu, M. Chatterjee, H. Schmid, S. Beck, and M. Gawaz, “Cxcl14 as an emerging immune and inflammatory modulator,” Journal of Inflammation, vol. 13, pp. 1–8, 2016.

[28] U. Lee, Y. R. Nam, J. S. Ye, K. J. Lee, N. Kim, and C. H. Joo, “Cysteine-rich secretory protein 3 inhibits hepatitis c virus at the initial phase of infection,” Biochemical and biophysical research communications, vol. 450, no. 2, pp. 1076–1082, 2014.

[29] D. C. Blaydon et al., “Mutations in csta, encoding cystatin a, underlie exfoliative ichthyosis and reveal a role for this protease inhibitor in cell-cell adhesion,” The American Journal of Human Genetics, vol. 89, no. 4, pp. 564–571, 2011.

[30] G. Wang, X. Zhang, J.-S. Lee, X. Wang, Z.-Q. Yang, and K. Zhang, “Endoplasmic reticulum factor erlin2 regulates cytosolic lipid content in cancer cells,” Biochemical Journal, vol. 446, no. 3, pp. 415– 425, 2012.

[31] S. Zhang et al., “Distinct prognostic values of s100 mrna expression in breast cancer,” Scientific reports, vol. 7, no. 1, p. 39 786, 2017.

[32] S. Lamouille, J. Xu, and R. Derynck, “Molecular mechanisms of epithelial–mesenchymal transition,” Nature reviews Molecular cell biology, vol. 15, no. 3, pp. 178–196, 2014.

[33] T. Chen, Y. You, H. Jiang, and Z. Z. Wang, “Epithelial–mesenchymal transition (emt): A biological process in the development, stem cell differentiation, and tumorigenesis,” Journal of cellular physiology, vol. 232, no. 12, pp. 3261–3272, 2017.

[34] R. P. Rastogi, A. Kumar, M. B. Tyagi, R. P. Sinha, et al., “Molecular mechanisms of ultraviolet radiation-induced dna damage and repair,” Journal of nucleic acids, vol. 2010, 2010.

[35] I. Kim and Y.-Y. He, “Ultraviolet radiation-induced non-melanoma skin cancer: Regulation of dna damage repair and inflammation,” Genes & diseases, vol. 1, no. 2, pp. 188–198, 2014.

[36] S.-W. Lin et al., “Prospective study of ultraviolet radiation exposure and risk of cancer in the united states,” International journal of cancer, vol. 131, no. 6, E1015–E1023, 2012.

[37] G. Nash, D. Walsh, and A. Kakkar, “The role of the coagulation system in tumour angiogenesis,” The lancet oncology, vol. 2, no. 10, pp. 608–613, 2001.

[38] F. Belardelli, “Role of interferons and other cytokines in the regulation of the immune response,” Apmis, vol. 103, no. 1-6, pp. 161–179, 1995.

[39] K. Schroder, P. J. Hertzog, T. Ravasi, and D. A. Hume, “Interferon-γ: An overview of signals, mechanisms and functions,” Journal of Leucocyte Biology, vol. 75, no. 2, pp. 163–189, 2004.

[40] A. Takaoka et al., “Integration of interferon-α/β signalling to p53 responses in tumour suppression and antiviral defence,” Nature, vol. 424, no. 6948, pp. 516–523, 2003.

[41] M. J. Walter et al., “Acquired copy number alterations in adult acute myeloid leukemia genomes,” Proceedings of the National Academy of Sciences, vol. 106, no. 31, pp. 12 950–12 955, 2009.

[42] C. Engen et al., “Flt3-itd mutations in acute myeloid leukaemia–molecular characteristics, distribution and numerical variation,” Molecular Oncology, vol. 15, no. 9, pp. 2300–2317, 2021.

[43] M. D. Wilkerson and D. N. Hayes, “Consensusclusterplus: A class discovery tool with confidence assessments and item tracking,” Bioinformatics, vol. 26, no. 12, pp. 1572–1573, 2010.

[44] H. Song et al., “Single-cell analysis of human primary prostate cancer reveals the heterogeneity of tumor-associated epithelial cell states,” Nature communications, vol. 13, no. 1, p. 141, 2022.

[45] K. P. Schliep, “Phangorn: Phylogenetic analysis in r,” Bioinformatics, vol. 27, no. 4, pp. 592–593, 2011.

[46] K. Street et al., “Slingshot: Cell lineage and pseudotime inference for single-cell transcriptomics,” BMC genomics, vol. 19, pp. 1–16, 2018.

[47] T. Wu et al., “Clusterprofiler 4.0: A universal enrichment tool for interpreting omics data,” The innovation, vol. 2, no. 3, 2021.

