## Supplementary material for "TUSCAN: Tumor segmentation and classification analysis in spatial transcriptomics": Supplementary files description.docx

**Table S1. Sources of Published Spatial Transcriptomics Datasets Used in This Study**

**Table S2: File used to construct tumor clone tree of human breast cancer data (block A section 1) clone 1 versus 2. This file summarizes the CNV state of each clone at different chromosome positions. The row names are all the chromosome bands where CNV appears, and the column names are clone A-F. We define five CNV states: 2-more gain, 1-less gain, 0-neutral, -1- less loss, -2-more loss.**

**Table S3: Chromosomal locations of copy number gains and losses of tumor clone A to F from human breast cancer (block A section 1) patient.**

**Table S4: Differentially expressed genes from human breast cancer data (block A section 1) clone 1 versus 2.**

**Table S5: Differentially expressed genes from human breast cancer data (block A section 1) clone 1 versus 3.**

**Table S6: Differentially expressed genes from human breast cancer data (block A section 1) clone 2 versus 3.**
