## Supplementary material for "TUSCAN: Tumor segmentation and classification analysis in spatial transcriptomics": Table S1.docx

**Supplementary Materials**

**Table S1. Sources of Published Spatial Transcriptomics Datasets Used in This Study**

| **Data** | **Sequencing Platform** | **Data Source** | **Data Summary** |
| --- | --- | --- | --- |
| **Human Breast Cancer: Ductal Carcinoma in Situ** | **10x Visium** | **10x Genomics**  [**https://www.10xgenomics.com/datasets/human-breast-cancer-ductal-carcinoma-in-situ-invasive-carcinoma-ffpe-1-standard-1-3-0**](https://www.10xgenomics.com/datasets/human-breast-cancer-ductal-carcinoma-in-situ-invasive-carcinoma-ffpe-1-standard-1-3-0) | **2518 spots, 17943 genes** |
| **Human breast cancer (Block A Section 1)** | **10x Visium** | **10x Genomics**  [**https://www.10xgenomics.com/datasets/human-breast-cancer-block-a-section-1-1-standard-1-1-0**](https://www.10xgenomics.com/datasets/human-breast-cancer-block-a-section-1-1-standard-1-1-0) | **3798 spots, 36,601 genes** |
| **Human prostate cancer adenocarcinoma with invasive carcinoma** | **10x Visium** | **10x Genomics**  [**https://www.10xgenomics.com/resources/datasets/human-prostate-cancer-adenocarcinoma-with-invasive-carcinoma-ffpe-1-standard-1-3-0**](https://www.10xgenomics.com/resources/datasets/human-prostate-cancer-adenocarcinoma-with-invasive-carcinoma-ffpe-1-standard-1-3-0) | **4371 spots, 17943 genes** |
| **Human HER2+ breast cancer** | **ST** | **Andersson, A., *et al*.**  [**https://github.com/almaan/her2st**](https://github.com/almaan/her2st) | **293 spots, 15109 genes** |
